# Improved Metagenomic Binning with Transformers

**DOI:** 10.1101/2022.02.12.479459

**Authors:** Nina Shenker-Tauris, Jeanette Gehrig

**Affiliations:** Technical University of Denmark (DTU); Siolta Therapeutics

## Abstract

Traditional metagenome binning methods cluster contiguous DNA sequences (contigs) based on uncontextualized features of the sequences which ignores both the semantic relationship between genes and the positional embedding of k-mers. This paper presents a novel binning method that addresses these concerns. Firstly, taken from natural language processing literature, a sequence representation model - Bidirectional Encoder Representations from Transformers (BERT) - is utilized to generate semantic and positional contig embeddings. Secondly, two workflows are presented; one which applies a hierarchical density-based clustering algorithm to find metagenomic bins and the other which incorporates contig embedding into a state-of-the-art binner. Experimental results on a publicly available metagenomic dataset show superior clustering for shorter contigs compared to traditionally used tetranucleotide frequency (TNF), reconstruction of up to 17% more high-precision genomes, and improved semantic understanding of contigs.

## 1 Introduction

Metagenomics is the study of microbial communities directly in their natural environments, bypassing the need for isolation and lab cultivation of individual species [1]. Metagenomics can be broken into two words “meta” and “genomics”. *Genomics* refers to analysis of genetic material and *meta* implies this analysis is applied to many organisms at once.

Metagenomics has been used to study both free-living and host-associated microbial communities. Of its many applications, metagenomics has been particularly useful in the study of the human microbiome. The human microbiota consists of trillions of microbial cells across many body sites [2] and the genomes and genes it harbors (the microbiome) have been showed to play a key role in both health and disease [3]. Dysbiosis of the microbiota has been linked to a variety of diseases such as asthma, allergies, type 2 diabetes, periodontitis, bacterial vaginosis, and inflammatory bowel disease [3–5].

As much of the human microbiome is anaerobic and difficult to culture, metagenomic data provides a key resource for assembling and obtaining microbial genomes [6]. In this paper, we tackle the task of binning, a vital step in metagenomic assembly. Binning clusters metagenomic sequences by organism of origin, enabling the reconstruction of known and unknown genomes from metagenomic sequences. Each sequence is placed into an imaginary bin representing ideally only fragments belonging to this group. A microbial community is usually a complex combination of multiple organisms and recovering their genomes is crucial to understand the behavior and functions within such communities [7]. Binning is key in profiling the microbiome and serves as a preliminary step in diagnosing patients, treating disease, and further exploration of the microbiome [3].

## 2 Methods

Just as words were converted to a numerical vector in the original BERT model [8], so is every contig-derived k-mer. To begin, since metagenomic contigs are often much longer than 512 k-mers (contigs may be up to millions of base pairs), the data must be pre-processed. Contigs are typically in FASTA format, with one FASTA file per microbiome sample. In FASTA files of contigs, each contig sequence is on a separate line, preceded by a header line containing a unique contig identifier.

As the first step of contig processing, the sequences are converted to k-mers. Any k-mer length would be acceptable, but k-mers of length *k* = 4^1^ are the standard for metagenomic analysis. K-mer representation is used as it creates a deeper contextual understanding of the sequence rather than regarding each base individually [9].

As the second step, the k-merized contigs are divided into segments of 512. For example, a k-merized contig of length 1024 is divided into two 512-segments. If a sequence length is not evenly divisible by 512, the remaining shorter segment is padded using the [PAD] token (Table 1).

**Table 1:**
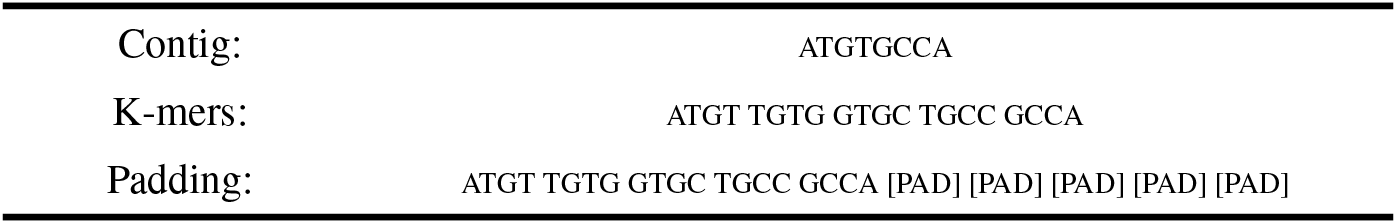
Example of pre-processing for contigs: first the contig is turned into k-mers and then padded to the desired length.

Here is a simplified example given the desired output of a contig of length 10:

### 2.1 Embedding of k-mers

When creating 4-mers from DNA there are 4^4^ or 256 possibilities of different base combinations. Given a vocabulary list, each unique k-mer corresponds to a numerical vector. For example, the k-mer aaaa is represented as the numerical vector “[5]”. Since the self-attention mechanism is not aware of token position, in order to provide the model with positional information, a position vector is assigned to every token. The sum of these embeddings are the final input into the model.

**Table 2:**
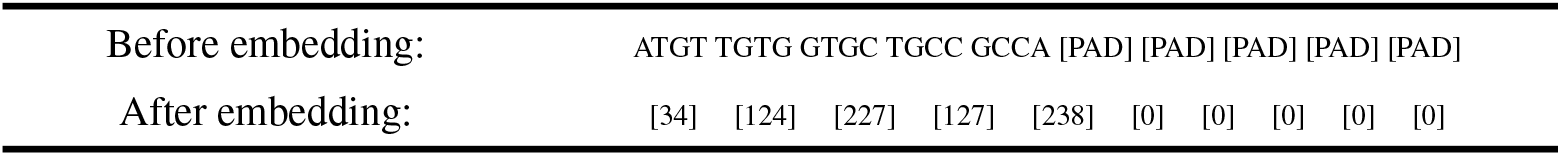
Example of embedding of k-mers: Every 4-mer combination is represented by a unique numerical representation.

### 2.2 Training tasks: modified Masked LM and NSP

The original BERT model was trained for several language tasks including sentence pair classification, single sentence classification, and question and answer task. Likewise, DNABERT [9] was trained for several genomic tasks including predicting gene promoters, identifying transcription factor binding sites, and predicting splice sites (Figure 1). Please refer to the BERT and DNABERT paper for further details [8, 9].

**Figure 1:**
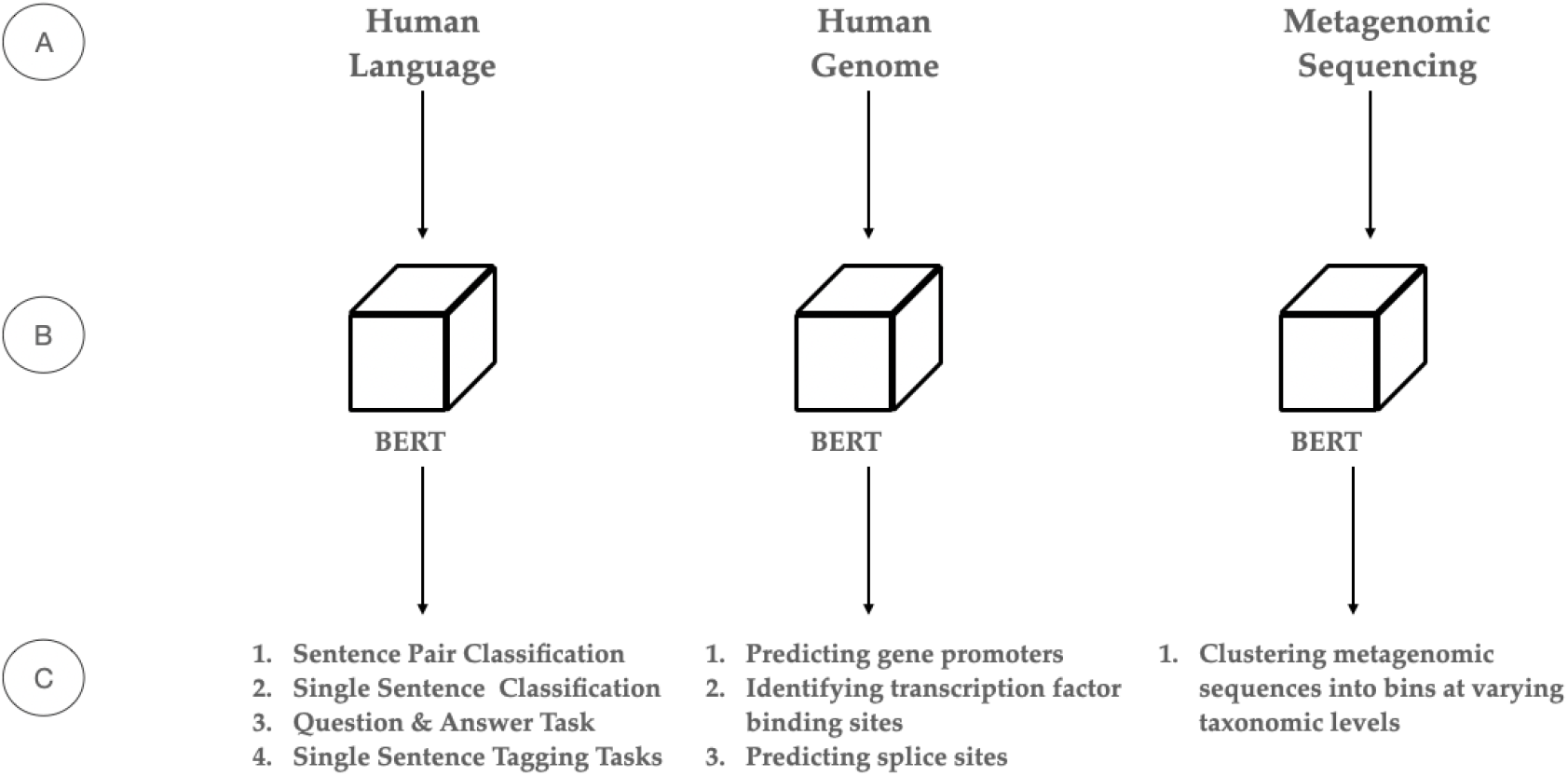
A) Type of data model is trained on, B) The BERT model, C) Downstream tasks for model

Training a BERT model is very resource-intensive. Therefore, we utilized the open-source DNABERT-4 model, as the base model for training metagenomic contigs. The DNABERT-4 base model was trained on the random segments of the human genome (240M sequences) for about 25 days on 8 NVIDIA 2080 Ti GPUs.

### 2.3 Training on metagenomic sequence data

Below are descriptions of the modified Masked LM and NSP task for training on metagenomics data.

#### Masked K-mers Task

While in human languages words in a sentence are discrete and meaning is derived from combining words, with DNA, each individual k-mer can be assumed based on the preceding and following k-mers, which makes masking difficult [9]. For example, a 4-mer sequence catg, atga, tgac, gact, “tgac” is the concatenation of “tga” in “atga” and “c” in “gact”. If 15% of k-mers are independently masked, in most instances, the previous and next k-mers of a masked one are unmasked. This will significantly simplify the pre-training task and prevent the model from learning deep semantic relations for a contig, as the masked token can be trivially inferred from the immediate adjacent tokens [9]. Therefore, instead of independently masking k-mers, we mask a contiguous span of k-mers. Here is a simplified example:

- The [MASK] token 80% of the time → e.g. atgg **tggt ggtt** gttattac → atgg **[mask] [mask]** gttattac
- A random k-mer 10% of the time → e.g. atgg **tggt ggtt** gtta ttac → atgg **catt gtac** gtta ttac
- An unchanged k-mer 10% of the time → e.g. atgg **tggt ggtt** gtta ttac → atgg **tggt ggtt** gtta ttac

At each step, a cross-entropy loss is calculated over all the masked k-mers.

#### Next Segment Prediction

The next segment prediction task is a modified version of the NSP task from the BERT. It was implemented to allow the model to get a better understanding of segments from the same contig. It consists of taking two masked segments, segment *A* and segment *B*, that are combined with the use of the [SEP] token. 50% of the time segment *B* is the next segment in the contig and the other 50% it is a random segment from another contig. The two segments get a combined label of “IsNext” or “NotNext”. For example, given two segments there are two possibilities:

- Input: **[cls]** atgg [mask] ggtt gtta **[sep]** tttg ttga [mask] gaac **[sep]** → Label = IsNext or 1
- Input: **[cls]** atgg [mask] ggtt gtta **[sep]** cggt ggta [mask] taac **[sep]** → Label = NotNext or 0

The NSP task is added on top of the masked k-mers task. Therefore, at every step the loss is defined as the combined cross-entropy loss for the masked k-mers and NSP task.

#### 2.3.1 Pooling of hidden layers

A majority of DNA applications can be categorized as sequence-level tasks and token-level tasks [9]. Metagenomic clustering is a sequence-level task, since entire contigs are binned, not individual k-mers. Thus, by feeding the embeddings to a BERT model, we obtain a matrix of the final contig segment representation.

For sequence-level tasks, the vector corresponding to the [CLS] token is fed to the output layer. The NSP task is considered a sequence level task while the masked k-mers task on its own is considered a token-level task as the goal is to predict individual tokens, not an overall sentence embedding. For token-level tasks, a [CLS] token is not obtained. Instead the vectors corresponding to all tokens are independently fed to the output layer.

In order to utilize the contig embedding, it is necessary to “pool” or “collapse” one of the 12 hidden layers. After the input passes through a hidden layer, each segment is translated to a N-dimensional vector, with N being the number of features embedded. To get a fixed representation of this, a dimensionality reduction is needed. This can be applying an aggregate function^2^, such as mean or max or if the [CLS] token is utilized, this can be used as it represents the entire contig not one token.

## 3 Workflows

The output of the BERT model can be used as an independent binner, we call **DeepBin**, or as an additional contig feature for the VAMB [10] workflow.

The output of the BERT model is a representation of the contig segments’ relationships to one another. Recalling an encoder, *E*, and its ability to encode features of data, we can represent *z* as an “encoded” version of the input, *x*. For every tokenized segment of length *L*, there are *N* features encoded 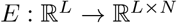 and thus:

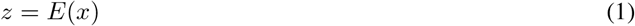

*z* is also called latent space, and will be referred to as such from now on. As data is embedded, the contig segments, each represented by a point in the latent space, can be imagined as coordinates in space where segments that are similar are closer together.

Due to the BERT model’s limited size input of 512, the latent space is not representative of the entire contig (if over 512 k-mers in length), but rather a 512-segment of the contig.

In the next section, we will introduce how to extract bins directly from the latent space, followed by the workflow of adding the latent space embeddings to the traditional VAMB workflow.

### 3.1 Overview of DeepBin binner

DeepBin’s workflow consists of two major steps: (1) forming bins from the latent space using a clustering algorithm and (2) evaluating the clusters using well-known clustering evaluation metrics.

#### 3.1.1 Clustering

In order to turn the latent space into bins, three clustering methods were tested. The clustering methods were:

1. K-Means[11]
2. K-Medoids [12]
3. HBDSCAN: Hierarchical Density-Based Spatial Clustering of Applications with Noise [13]

Each method was implemented by taking the latent space projection, represented by a tensor^3^ and clustering with each method’s unique criteria. All clustering algorithms were run with default settings and tested using a variety of cluster sizes (Table 3).

**Table 3:** Settings of each clustering method; n = number of genomes

| Clustering Algorithm | Cluster Size | Contributor |
| --- | --- | --- |
| K-Means | 0.2*n, 0.5*n, n | scikit-learn [14] |
| K-Medoids | 0.2*n, 0.5*n, n | scikit-learn [14] |
| HDBSCAN | 5, 10, 15, 20 | scikit-learn-contrib [15] |

#### 3.1.2 Evaluation of DeepBin

Evaluation of DeepBin’s clustering involves comparison to a ground truth reference file. To compare bins, the following evaluation metrics were chosen: recall, precision, accuracy, and F1 score.

All metrics were reported as averages and higher is better. Small bins contribute in the same way as large bins. Specifically, the average precision is the fraction of correctly assigned base pairs for all assignments to a given bin averaged over all predicted genome bins, where unmapped genomes are not considered. This value reflects how trustworthy the bin assignments are on average. The recall is averaged over all genomes, including those not mapped to genome bins (for which recall is zero) [16]. In addition, Rand Index, a measure of similarity, was calculated when benchmarking DeepBin and tetranucelotide frequency (TNF) against a reference. These metrics are commonly used for clustering and have been used in previous metagenomic benchmarking literature [17, 18].

Firstly, a 2×2 similarity matrix was generated using sci-kit-learn’s pair confusion matrix [14]. This matrix was created by comparing the clusters computed by DeepBin to the ground truth clusters computed from a reference file. The similarity matrix contains the following entries:

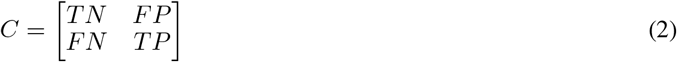

- TN: True Negatives, the number of pairs with both clusterings having the samples not clustered together.
- FP: False Positives, the number of pairs with the ground truth clustering not having the samples clustered together, but DeepBin’s clustering having the samples clustered together.
- FN: False Negatives, the number of pairs with the ground truth clustering having the samples clustered together, but DeepBin’s clustering does not.
- TP: True Positives, the number of pairs where both clusterings have the samples clustered together.

All of the clustering metrics can be calculated from the confusion matrix. The following equations were used to calculate the specified metrics:

**Recall:**

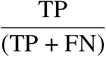
**Precision:**

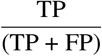
**Rand Index/Accuracy^4^:**

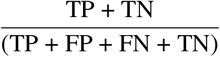
**F1 Score^5^:**

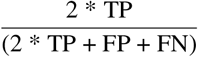

### 3.2 Input into VAMB

The output of the BERT model can be used in addition to TNF and abundance for the VAMB [10] workflow. Below we present a brief overview of the VAMB workflow with contig embedding. For a more detailed explanation, please refer to “Improved metagenome binning and assembly using deep variational autoencoders” [10].

VAMB workflow with contig embedding (additions to the original workflow are bolded):

1. The input into the VAMB pipeline is (a) metagenomic sequences (FASTA files), (b) abundance information (FASTQ files), and (c) **contig embedding**.
2. For every contig, TNF is calculated from a FASTA file and abundance is calculated from BAM files^6^. **As described above, the output of the BERT model produces a latent space embedding for a 512-segment from every contig.** This means while TNF and abundance are calculated from the entire contig sequence, **the contig embedding is only representative of a 512-segment**.
3. TNF, abundance, and **contig embedding** are all represented by their respective vector: TNF vector of size 103, abundance vector of size *n* or number of samples, and **contig embedding of size 768^7^**.
4. The three features TNF, abundance, **contig embedding** are concatenated and used to train a variational autoencoder (VAE).
5. Following training, the features are encoded by the mean of their latent space distribution.
6. VAMB’s iterative medoid clustering algorithm is applied.

The workflows were evaluated using VAMB’s evaluation code. Here, the true positives, true negatives, false positives, and false negatives are calculated for each bin. To determine if the contig embedding information lead to better binning by the VAE, recall-precision metrics were calculated for three taxonomic levels: genus, species, and strain. The VAMB creators chose near-complete (Nc) bins as their evaluation metric. This means the recall and precision metrics were set to 0.9 and 0.95 respectively. Only bins that met or exceeded the precision and recall thresholds in comparison to ground truth genomes were considered “recovered” [10].

**Figure 2:**
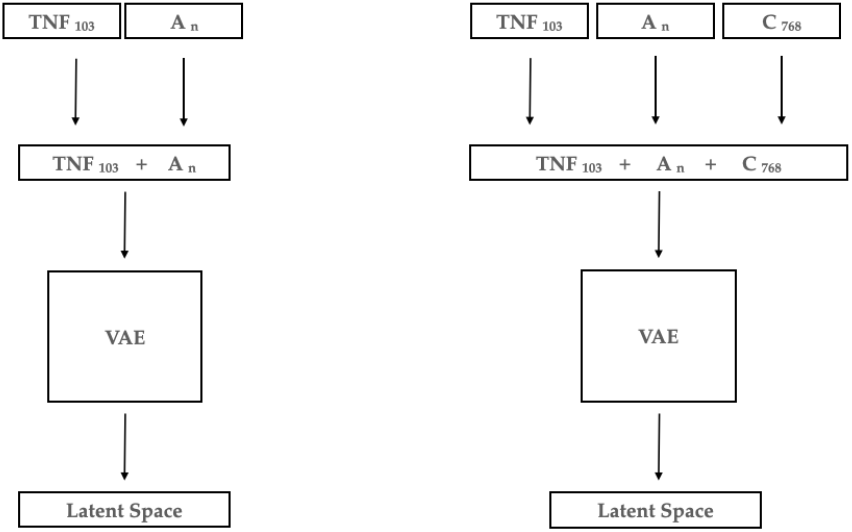
On left: traditional VAMB workflow; On right: modified VAMB workflow with contig embedding; A = abundance, C = contig embedding

#### 3.2.1 Loss function

Here, we present an overview of the new loss function.

When training a VAE, the loss is calculated by the failure to reconstruct the input or the reconstruction error. The error for contig embedding, *E_c_*, was defined as:

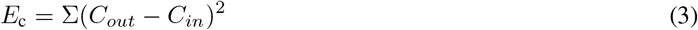

Using sum of squared errors (SSE) to represent the contig embedding loss, this loss is combined with the cross-entropy (CE) loss for abundance and SSE loss for TNF defined in VAMB’s methods.

The combined model loss, *L*, was thus:

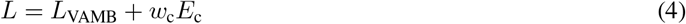

with *L*_VAMB_ being the loss function before the addition of contig embedding. The weighting term was defined as *w_c_* = *α*/768 with the parameter of *α* being set by the VAMB authors to 0.15.

## 4 Results

### Data access and preprocessing

The results presented in this chapter are based on benchmarking of two out of five synthetic datasets from the *2nd Critical Assessment of Metagenome Interpretation (CAMI) Toy Human Microbiome Project Datasets* [17]. The two datasets were the CAMI Oral, representing the human oral microbiota, and CAMI Airway, representing the human airway microbiota. Both datasets used in this study are publicly available and can be downloaded from the CAMI portal at data.cami-challenge.org. The synthetic data is simulated Illumina HiSeq metagenomic data with 2×150 basepairs (bp) read length and 270 bp insert size.

### Data partition

The CAMI Oral dataset was used for training the Transformer model. The training data of 10 samples was split into a 90/10 training split with the model training on nine samples and validating on one. For further experiments, only the nine train samples were used. We optimize training with AdamW and drop-out, using the same values as for the BERT modeld. The adjusted hyper-parameters that differ from the original BERT model are reported in Table 4. No ablation studies were done for hyper-parameter tuning. The training and validation samples may be used to fit model parameters or to determine hyperparameter values. The CAMI Airway dataset (10 samples) was used as test data. The test dataset is used to observe if the model is able to generalize on unseen data. More information on the datasets are found in the appendix (Table 15).

**Table 4:** Hyperparameters for training of DeepBin

| Hyper-Parameter |  |
| --- | --- |
| Batch Size | 8 |
| Learning Rate | 2e-5 |
| Epochs (Masking only) | 20 |
| Epochs (Masking + NSP) | 40 |

**Table 5:** Evaluation metrics for clustering the latent space at species level

| Clustering Method | Cluster Size | Recall | Precision |
| --- | --- | --- | --- |
| K-Means | 5 | 0.56 | 0.56 |
| K-Means | 14 | 0.69 | 0.26 |
| K-Means | 28 | 0.71 | 0.13 |
| K-Medoids | 5 | 0.56 | 0.59 |
| K-Medoids | 14 | 0.61 | 0.23 |
| K-Medoids | 28 | **0.81** | 0.19 |
| HDBSCAN | 5 | 0.38 | 0.71 |
| HDBSCAN | 10 | 0.22 | 0.93 |
| HDBSCAN | 15 | 0.22 | 0.94 |
| HDBSCAN | 20 | 0.22 | **0.95** |

**Table 6:** Evaluation of BERT after 20 epochs of masking training and after an additional 20 epochs of masking + NSP training; NSP = next segment prediction

| Training Task | Recall | Precision | Accuracy | F1 Score |
| --- | --- | --- | --- | --- |
| Masked k-mer only | 0.22 | 0.99 | 0.23 | 0.36 |
| Masked k-mer and NSP | 0.22 | 0.99 | 0.23 | 0.36 |

**Table 7:**
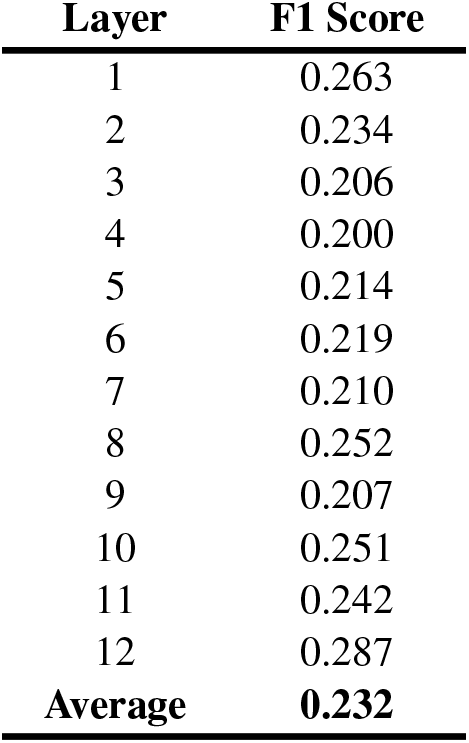
K-medoids clustering: Clustering F1 score for every hidden layer of the highest recall method, K-medoids

| **Layer** | **F1 Score** |
| --- | --- |
| 1 | 0.263 |
| 2 | 0.234 |
| 3 | 0.206 |
| 4 | 0.200 |
| 5 | 0.214 |
| 6 | 0.219 |
| 7 | 0.210 |
| 8 | 0.252 |
| 9 | 0.207 |
| 10 | 0.251 |
| 11 | 0.242 |
| 12 | 0.287 |
| **Average** | **0.232** |

**Table 8:** HDBSCAN clustering: Clustering F1 score for every hidden layer of the highest precision method, HDBSCAN

| **Layer** | **F1 Score** |
| --- | --- |
| 1 | 0.516 |
| 2 | 0.489 |
| 3 | 0.537 |
| 4 | 0.533 |
| 5 | 0.513 |
| 6 | 0.517 |
| 7 | 0.681 |
| 8 | 0.364 |
| 9 | 0.359 |
| 10 | 0.358 |
| 11 | 0.358 |
| 12 | 0.361 |
| **Average** | **0.465** |

**Table 9:** Oral dataset; Number of bins at **strain** level reconstructed with a precision of at least 95%. Number of additional bins the contig embedding component provided are highlighted for each recall level.

| Binner | Recall |  |  |  |  |  |  |  |  |
| --- | --- | --- | --- | --- | --- | --- | --- | --- | --- |
|  | 0.30 | 0.40 | 0.50 | 0.60 | 0.70 | 0.80 | 0.90 | 0.95 | 0.99 |
| VAMB | 178 | 162 | 153 | 142 | 137 | 125 | 115 | 97 | 80 |
| VAMB + Contig embedding | 188 | 175 | 161 | 153 | 147 | 136 | 116 | 98 | 80 |
| Difference | **10** | **13** | **8** | **11** | **10** | **11** | **1** | **1** | **0** |

**Table 10:** Oral dataset; Number of bins at **species** level reconstructed with a precision of at least 95%. Number of additional bins the contig embedding component provided are highlighted for each recall level.

| Binner | Recall |  |  |  |  |  |  |  |  |
| --- | --- | --- | --- | --- | --- | --- | --- | --- | --- |
|  | 0.30 | 0.40 | 0.50 | 0.60 | 0.70 | 0.80 | 0.90 | 0.95 | 0.99 |
| VAMB | 117 | 110 | 105 | 100 | 97 | 92 | 84 | 71 | 58 |
| VAMB + Contig embedding | 128 | 122 | 116 | 112 | 109 | 103 | 87 | 74 | 60 |
| Difference | **11** | **12** | **11** | **12** | **12** | **11** | **3** | **3** | **2** |

**Table 11:** Oral dataset; Number of bins at **genus** level reconstructed with a precision of at least 95%. Number of additional bins the contig embedding component provided are highlighted for each recall level.

| Binner | Recall |  |  |  |  |  |  |  |  |
| --- | --- | --- | --- | --- | --- | --- | --- | --- | --- |
|  | 0.30 | 0.40 | 0.50 | 0.60 | 0.70 | 0.80 | 0.90 | 0.95 | 0.99 |
| VAMB | 58 | 57 | 54 | 52 | 51 | 47 | 46 | 41 | 34 |
| VAMB + Contig embedding | 65 | 63 | 60 | 58 | 58 | 55 | 50 | 42 | 36 |
| Difference | **7** | **6** | **6** | **6** | **7** | **8** | **4** | **1** | **2** |

**Table 12:** Airway dataset; Number of bins at **strain** level reconstructed with a precision of at least 95%. The difference in the total number of bins reconstructed for each method was highlighted for each recall level.

| Binner | Recall |  |  |  |  |  |  |  |  |
| --- | --- | --- | --- | --- | --- | --- | --- | --- | --- |
|  | 0.30 | 0.40 | 0.50 | 0.60 | 0.70 | 0.80 | 0.90 | 0.95 | 0.99 |
| VAMB | 130 | 115 | 111 | 100 | 87 | 74 | 59 | 50 | 39 |
| VAMB + Contig embedding | 129 | 115 | 111 | 103 | 87 | 78 | 59 | 50 | 41 |
| Difference | **-1** | **0** | **0** | **3** | **0** | **4** | **0** | **0** | **2** |

**Table 13:**
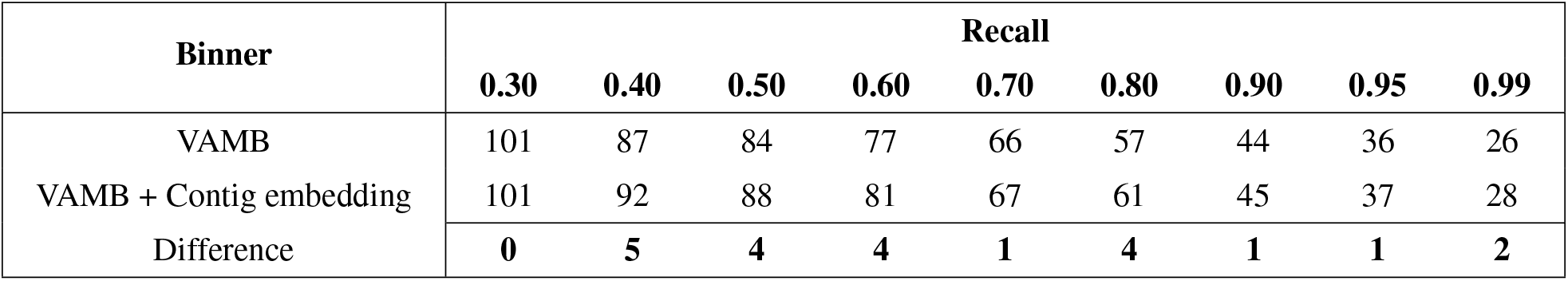
Airway dataset; Number of bins at **species** level reconstructed with a precision of at least 95%. The difference in the total number of bins reconstructed for each method was highlighted for each recall level.

| Binner | Recall |  |  |  |  |  |  |  |  |
| --- | --- | --- | --- | --- | --- | --- | --- | --- | --- |
|  | 0.30 | 0.40 | 0.50 | 0.60 | 0.70 | 0.80 | 0.90 | 0.95 | 0.99 |
| VAMB | 101 | 87 | 84 | 77 | 66 | 57 | 44 | 36 | 26 |
| VAMB + Contig embedding | 101 | 92 | 88 | 81 | 67 | 61 | 45 | 37 | 28 |
| Difference | **0** | **5** | **4** | **4** | **1** | **4** | **1** | **1** | **2** |

**Table 14:** Airway dataset; Number of bins at **genus** level reconstructed with a precision of at least 95%. The difference in the total number of bins reconstructed for each method was highlighted for each recall level.

| Binner | Recall |  |  |  |  |  |  |  |  |
| --- | --- | --- | --- | --- | --- | --- | --- | --- | --- |
|  | 0.30 | 0.40 | 0.50 | 0.60 | 0.70 | 0.80 | 0.90 | 0.95 | 0.99 |
| VAMB | 52 | 44 | 43 | 38 | 33 | 30 | 20 | 13 | 9 |
| VAMB + Contig embedding | 52 | 47 | 44 | 39 | 33 | 30 | 21 | 13 | 9 |
| Difference | **0** | **3** | **1** | **1** | **0** | **0** | **1** | **0** | **0** |

**Table 15:** Summary statistics for datasets. Source: VAMB [10]; Top 10 abundant species in CAMI Oral: *Campy-lobacter jejuni*, *Neisseria meningitidis*, *Clostridioides difficile*, *Streptococcus pyogenes*, *Burkholderia pseudomallei*, *Streptococcus pneumoniae*, *Streptococcus suis*, *Streptococcus agalactiae*, *Lactococcus lactis*, *Pasteurella multocida*; Top 10 abundant species in CAMI Airway: *Staphylococcus aureus*, *Corynebacterium pseudotuberculosis*, *Neisseria meningitidis*, *Klebsiella pneumoniae*, *Corynebacterium glutamicum*, *Corynebacterium diphtheriae*, *Streptococcus pneumoniae*, *Streptococcus pyogenes*, *Escherichia coli*, Streptococcus agalactiae

| Dataset | Samples | Strains | Species | Genera | Contigs | Assembly size | N50 | $H$ |
| --- | --- | --- | --- | --- | --- | --- | --- | --- |
| CAMI Oral | 10 | 673 | 249 | 107 | 201,563 | 1,938,340,074 | 28,290 | 4.39 |
| CAMI Airways | 10 | 639 | 311 | 139 | 187,685 | 1,653,971,669 | 25,427 | 3.89 |

The BERT model was trained for 20 epochs on the masked-kmers task followed by 20 epochs of the masked k-mers task combined with next segment prediction. It was run on a single NVIDIA V100 GPU (16 GB) through Amazon Web Services (AWS).

### Evaluation

Reference files for both the CAMI Oral and CAMI Airway datasets were created by combining the taxonomy and ground truth mapping files that map the contig name to the correct genome (provided by CAMI).

The BERT model was evaluated at inference. During inference, the model was only given positive examples, meaning unmasked segments and two connected contig segments. The model was tested from inference, after 20 and 40 epochs, by evaluating on a random subsample (8000 contigs) from the CAMI Oral Dataset. For evaluation, the shortest contigs without a “next segment” were filtered.

When evaluating the impact of adding contig embedding to VAMB, the workflow described in the paper [10] was run. For all datasets only contigs > 2, 000 base pairs (bp) were input and VAMB was run with default parameters and bin-splitting enabled.

For the following experiments, unless otherwise specified, the contig embedding was generated from the last hidden layer, collapsed by the mean of the 512-dimension, clustered with HDBSCAN (cluster size 20), and binned at species level.

### 4.1 Comparison against TNF

In order to compare DeepBin to TNF, we calculated the Rand Index score for each of their respective bins. To test if there was variation in signal depending on contig length we divided the contigs into categories based on their length. Additionally, to test if DeepBin performed equally on training data vs unseen data, we tested on one train sample and one validation sample from the CAMI Oral Dataset. It is important to note that TNF is calculated for the entire contig while DeepBin’s embedding is only for a 512-segment of the contig.

Results show a general trend of DeepBin having superior clustering for shorter contigs while TNF frequency is superior for the longest contigs (Figure 3).

**Figure 3:**
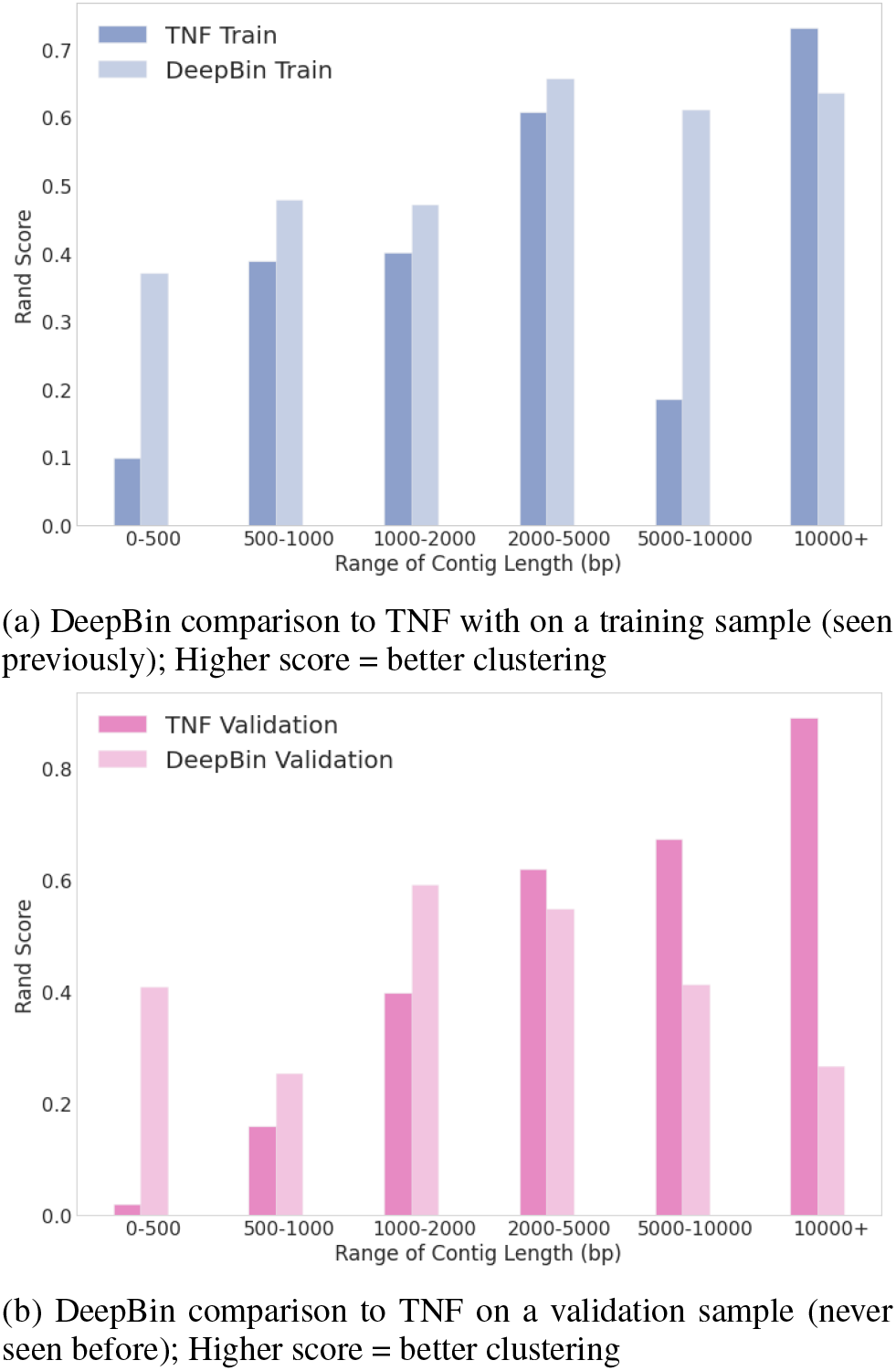
DeepBin comparison to TNF across categories based on contig length

DeepBin had a higher Rand Index for most groups when given training data vs unseen data. For the training data, DeepBin gets gradually better as contig length increases but is surpassed by TNF at contigs over 10,000 bp in length. For the validation data, DeepBin appears to have lower values than TNF from contigs length 2000 and above. Additionally, it appears to perform better for medium-length contigs (1000-5000) rather than improving as contigs get longer.

The results for TNF were mostly consistent across the train and validation sample. There was one deviation observed for the train dataset for contigs of length 5,000-10,000. The Rand index was 0.185 for the train sample, significantly lower than the 0.672 value reported for the validation sample.

### 4.2 Latent space visualization and clustering

Each hidden layer of the Transformer model consists of a 512 × 768 feature vector. In order to determine the best way to extract features from the neural network, each layer was “collapsed” or “pooled”. Pooling is necessary to get the entire sequence embedding rather than the individual k-mer embeddings. As mentioned in Section 2.3.1, for the NSP task a [CLS] token can be used for collapsing but for the masked k-mer task only, it can’t. Here we test after 20 epochs of masked k-mer training and thus require the layer to be collapsed. This was done by testing five collapse functions: max of the 768-dimension, max of the 512-dimension, mean of the 768-dimension, mean of the 512-dimension, and flattening of both layers by multiplying the 512-dimension by the 768-dimension.

From a training sample of the CAMI Oral dataset, 8000 contigs were randomly selected for the 10 most abundant species. The latent space embeddings were visualized (Figure 4) as a product of each of the five collapsing methods. Each point in the latent space represents a 512-segment embedding for a particular contig.

**Figure 4:**
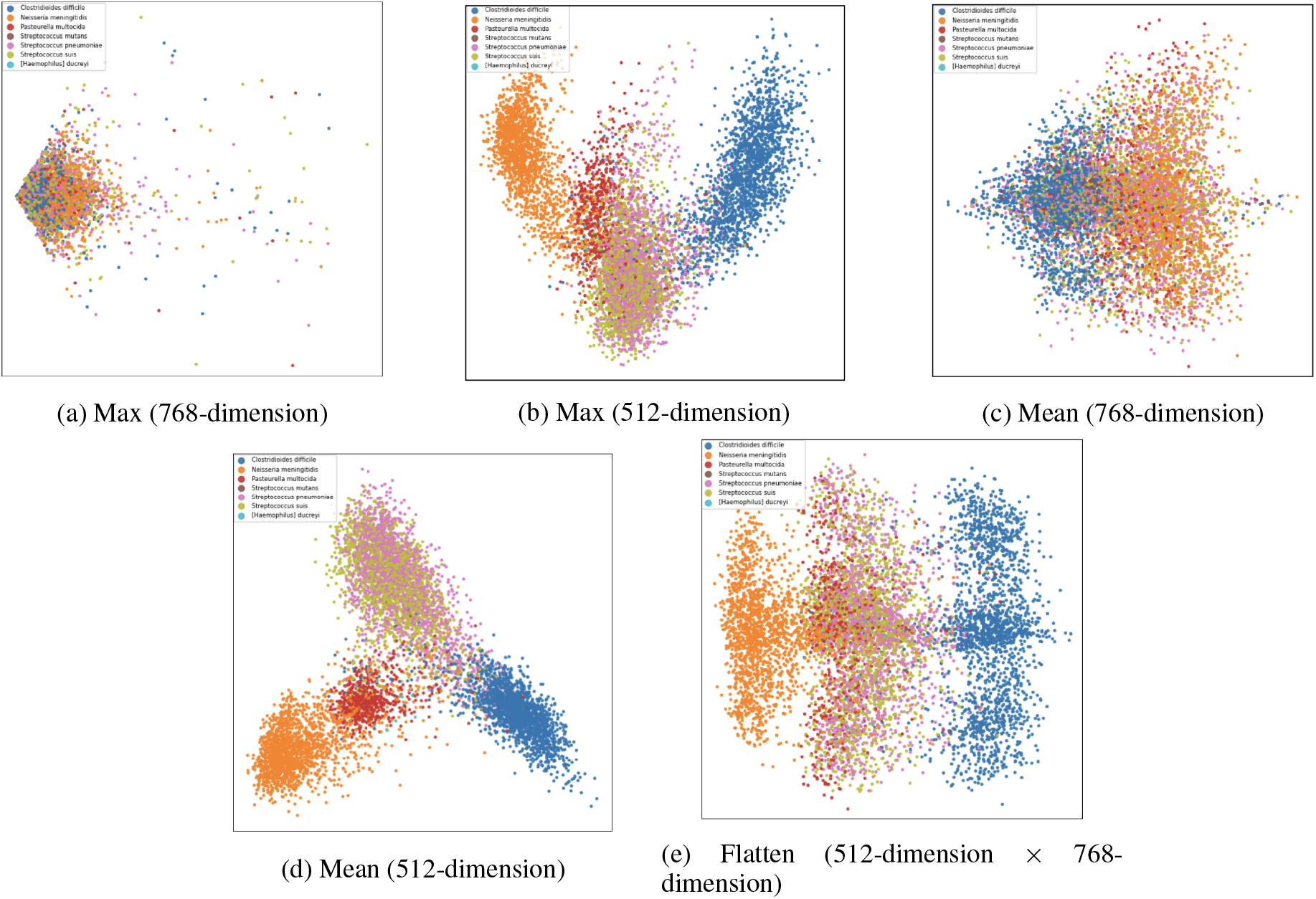
Latent space visualization of tested collapse functions for masking-only DeepBin model; *Blue*: *Clostridioides difficile*, *Orange*: *Neisseria meningitidis*, *Red*: *Pasteurella multocida*, *Brown*: *Streptococcus mutans*, *Pink*: *Streptococcus pneumoniae*, *Green*: *Streptococcus suis*, *Light blue*: *Haemophilus ducreyi*

Based on the visualizations, it appears the most clearly defined clusters were obtained taking the mean of the 512-dimension. These clusters were determined to be the most tightly bound while having the greatest separation from one another. Given these results, we use mean of the 512-dimension in the remainder of the experiments. However, it should be noted these visualizations are only representative of 10 genomes and therefore, may not be representative of the entire dataset or for binning at a different taxonomic level.

### 4.3 Clustering algorithms

An ablation study was performed to determine which clustering method was best suited for clustering the latent space. This was determined by visualizing the latent space of a single train sample along with clustering evaluation metrics. The three clustering methods; K-Means, K-Medoids, and HDBSCAN were compared and tested with multiple clustering sizes.

The clustering algorithms were also compared by visualizing the latent space clustering. The top ten most common species were visualized and clustered against the ground truth (Figure 7–10).

**Figure 5:**
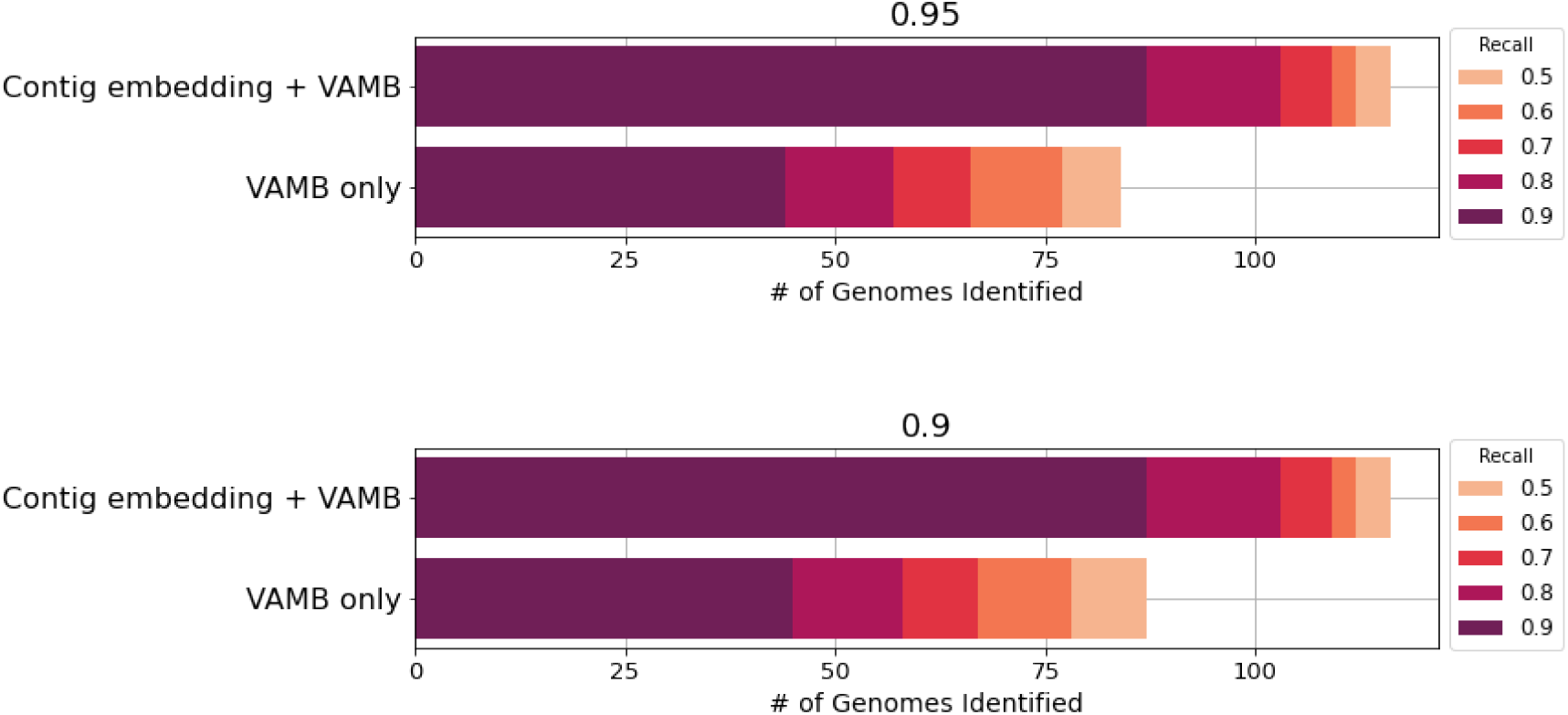
Oral dataset: High precision bin reconstructions by VAMB and Contig embedding + VAMB; Top: 95% precision, Bottom: 90% precision.

**Figure 6:**
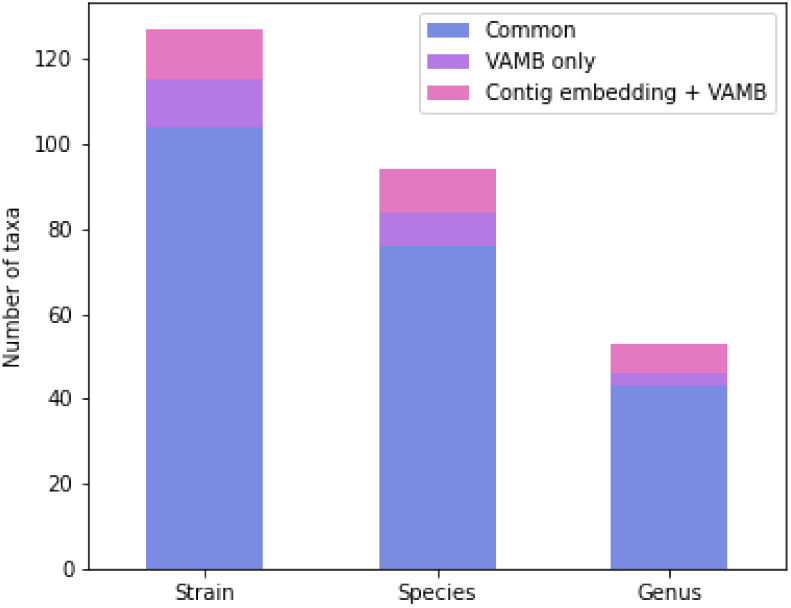
Number of taxa reconstructed with at least one NC bin by the two binners at three taxonomic levels; taxa identified by both methods (blue), taxa identified by VAMB only (purple) and VAMB + contig embedding (pink)

**Figure 7:**
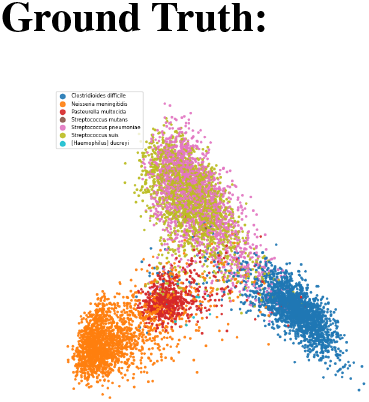
Ground Truth for clusters

**Figure 8:**
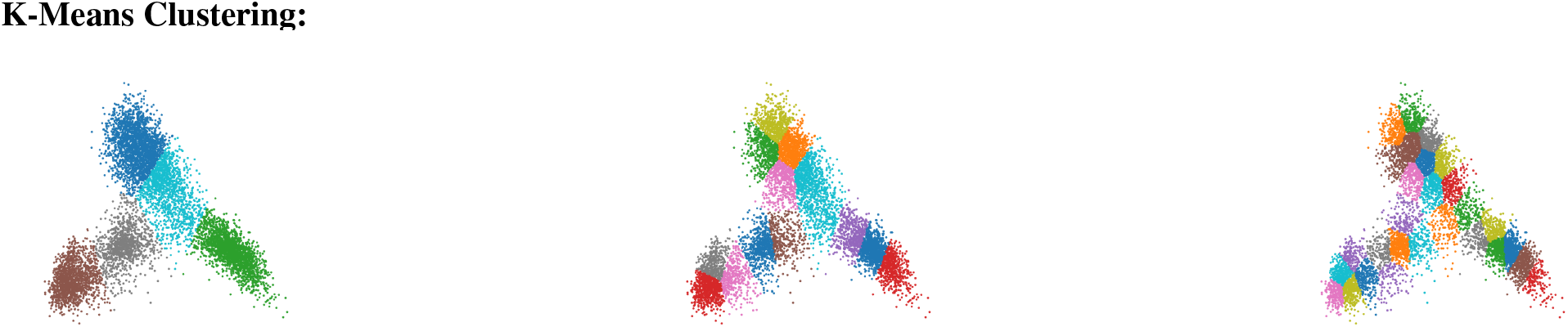
Cluster size is as follows: k=5, k=14, k=28 (left to right)

**Figure 9:**
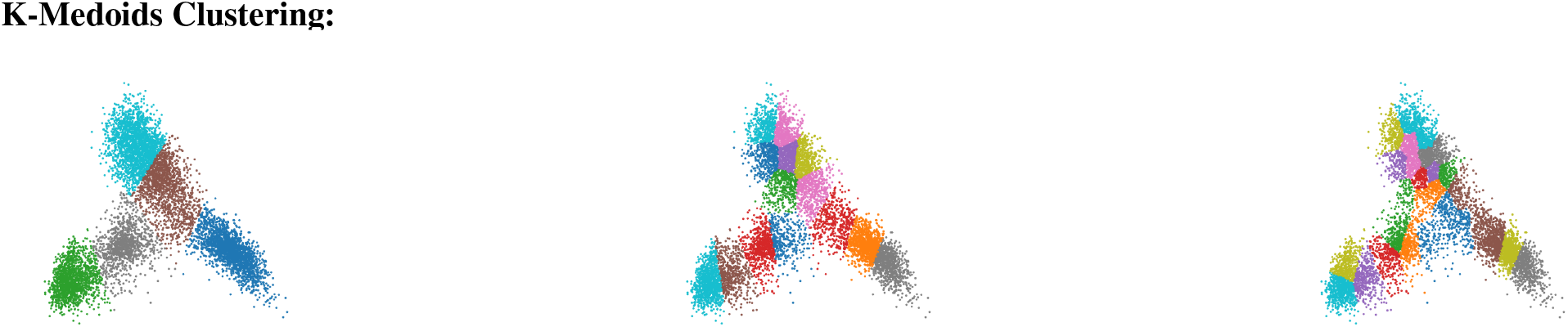
Cluster size is as follows: k=5, k=14, k=28 (left to right)

**Figure 10:**
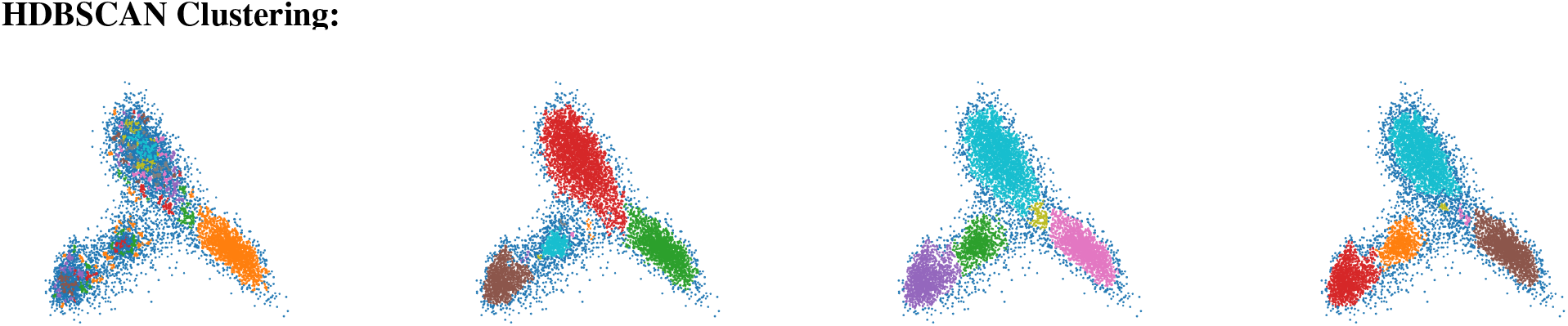
Cluster size is as follows: n=5, n=10, n=15, n=20 (left to right)

K-Means and K-Medoids clustering led to a high recall with low precision. There was a direct relationship between cluster size and recall; as cluster size increased, recall increased. Conversely, there was an inverse relationship between cluster size and precision. HDBSCAN had very high average precision for all cluster sizes over 10 but conversely had relatively low recall. The highest recall and precision method are bolded in Figure 5.

To further compare the highest recall (K-Medoids) and highest precision (HDBSCAN) clustering method, the F1 score (measurement that combines recall and precision) was computed for both methods at every hidden layer (12 total). Results show the average F1 score across all layers was for 0.465 for the HDBSCAN clustering method (Table 7) and 0.232 for the K-Medoids clustering method (Table 8).

Based on the higher average F1 score reported for HDBSCAN (cluster size 20), this clustering method was used for the remaining of the experiments. However, given the research task, one may opt for the K-Medoids clustering method if a higher recall is preferred to higher precision. The clustering results for this experiment are only indicative for a single randomly selected train sample and may not be representative of the entire CAMI Oral dataset or other datasets tested.

### 4.4 Addition of NSP

A comparison between the BERT model after 20 epochs of the masked k-mer task was compared to the same model after an additional 20 epochs of a masked k-mer + NSP combined task. The average recall and precision for CAMI Oral Dataset (training samples only) were recorded in order to see which model produced the best bins for species-level binning (Table 6).

The goal of combining the NSP task with the masked k-mers task was to simultaneously develop a deeper understanding of each contig segment while also improving the model’s knowledge of the relationships between segments of a contig.

Despite 20 epochs of additional training or 40 epochs of combined training, the BERT model showed equivalent results for all clustering metrics as it had done after the first 20 epochs.

### 4.5 VAMB with contig embedding

Results of training a VAE with the addition of contig embedding to TNF and abundance, showed an increase in the number of reconstructed bins at every taxonomic level (genus, species, strain) compared to a VAE trained only on TNF and abundance. The increase in reconstructed bins varied across taxonomic level with the greatest improvement being a 17% increase in genus level bins (0.8 recall or higher). The number of genomes reconstructed with 95% and 90% precision using VAMB and VAMB + contig embedding are shown for species level binning in Figure 5. Refer to Table 9, Table 10, and Table 11 for numerical results at strain, species, and genus level, respectively.

In order to determine, if contig embedding was able to generalize on unseen data, we additionally tested the workflows on the CAMI Airway Dataset. Results show the contig embedding workflow overall obtained more high-precision bins for the CAMI Airway dataset, as it had on the CAMI Oral dataset. Refer to Table 12, Table 13, and Table 14 for numerical results at strain, species, and genus level, respectively.

### 4.6 Further analysis of reconstructed bins

To establish overlapping bins between VAMB with and without contig embedding, we investigated the overlap of NC bins^8^ by the two methods for the CAMI Oral dataset (Figure 6). Unique taxa was discovered by mapping each bin to the corresponding genome/species/genus.

Investigating the taxonomic annotations showed a large overlap between the two set of bins. Additionally, results showed both binners had unique taxa for all three taxonomic levels. VAMB without contig embedding had 11 unique strains, 9 unique species, and 3 unique genera (Table 16). VAMB with contig embedding had 12 unique strains, 10 unique species, and 6 unique genera (Table 17). Thus combining the results of both binners, resulted in an overall increase for the number of NC reconstructed bins at all taxonomic levels. Further investigation is needed to determine why contig embedding improved binning for certain contigs while hindering binning for others.

**Table 16:** Unique taxa for VAMB only bins; Strains identified on NCBI Taxonomy [26]

| Taxonomic Level | Unique taxa |
| --- | --- |
| Genus | *Corynebacterium*, *Campylobacter*, *Mycoplasma* |
| Species | *Streptococcus pyogenes*, *Corynebacterium pseudotuberculosis*, *Streptococcus gallolyticus*, *Streptococcus pneumoniae*, *Treponema brennaborense*, *Burkholderia cepacia*, *Campylobacter coli*, *Treponema denticola*, *Mycoplasma capricolum* |
| Strain | Unclassified *Streptococcus pyogenes* strain, Unclassified *Corynebacterium pseudotuberculosis* strain, Unclassified *Streptococcus gallolyticus* strain, *Streptococcus pneumoniae* str. D39, Unclassified *Corynebacterium pseudotuberculosis* strain, *Treponema brennaborense* str. DSM 12168, *Burkholderia cepacia* str. ATCC 25416, Unclassified *Campylobacter coli* strain, Unclassified *Streptococcus agalactiae* strain, *Treponema denticola* str. ATCC 35405, *Mycoplasma capricolum* subsp. *capripneumoniae* str. M1601 |

**Table 17:** Unique taxa for VAMB + contig embedding bins; Strains identified on NCBI Taxonomy [26]

| Taxonomic Level | Unique taxa |
| --- | --- |
| Genus | *Riemerella*, *Myroides*, *Tenacibaculum*, *Maribacter*, *Ilyobacter*, *Neisseria* |
| Species | *Treponema putidum*, *Riemerella anatipestifer*, *Myroides profundus*, *Tenacibaculum* sp. LPB0136, *Burkholderia* sp. RPE64, *Burkholderia pseudomallei*, *Riemerella anatipestifer*, *Maribacter* sp. T28, *Ilyobacter polytropus*, *Neisseria meningitidis* |
| Strain | Unclassified *Treponema putidum* strain, Unclassified *Streptococcus agalactiae* strain, Unclassified *Riemerella anatipestifer* strain, Unclassified *Streptococcus agalactiae* strain, Unclassified *Myroides profundus* strain, Unclassified *Tenacibaculum* sp. LPB0136 strain, Unclassified *Burkholderia* sp. RPE64 strain, Unclassified *Burkholderia pseudomallei* strain, *Riemerella anatipestifer* str. RA-CH-1, Unclassified *Maribacter* sp. T28 strain, *Ilyobacter polytropus* str. DSM 2926, *Neisseria meningitidis* str. WUE 2594 |

## 5 Discussion

### 5.1 Contig embedding enables richer understanding and increases the number of reconstructed bins

At the beginning of this thesis we posed the research question of “*Can we input raw contigs into a state-of-the-art deep learning architecture and obtain better binning results?*”. The first part of this question can be answered with a simple “yes”. By training the BERT model on metagenomic sequencing, we was able to create a deep learning model that takes raw contigs as input. With this method, we created a novel contig embedding feature that is able to develop a more complex and holistic interpretation of a contig. The second part of this question is harder to answer than a simple “yes” or “no”. While we cannot conclude that contig embedding was able to obtain state-of-the-art results on its own, it brought focus to alternatives or additions to traditional binning features and methods. We showed how supplementing contig embedding with TNF and abundance for a state-of-the art method (VAMB) provides new state-of-the-art results. Contig embedding can be used for the reconstruction of more high precision bins at all three taxonomic levels for both trained and unseen data. Furthermore, comparing contig embedding to TNF, results showed contig embedding was consistent across all contig lengths compared to TNF that performed significantly poorer for short contigs.

### 5.2 Hand-crafted features vs learned features

Throughout this thesis, the topic of hand-crafted features vs learned feature representation, has been highlighted.

Looking at hand-crafted features vs. learned features, there are two key distinctions to consider. First, with a handcrafted feature such as TNF, we are simply looking at a small-scale pattern. There is no guarantee that several organisms don’t have the same TNF signal and furthermore, there is a risk of the same organism *not* having the same TNF signal across a genome. With contig embedding, we capture global contextual information from the entire input^9^.

While the first reasoning leans in contig embeddings favor, we must consider the computational expense and time. TNF has the advantage of being both fast and easy to calculate. Furthermore, when we analyze sequences based on this criteria we have a conceptual understanding of what is happening. When trying to conceptualize the BERT model, it becomes difficult. The BERT encodes 768 features for every contig, but what actually are these features? While we imagine one could be a variation of k-mer frequency and another could be a semantic understanding of which k-mers often follow each other, there is no way of knowing exactly what features the BERT model is creating. This is why deep learning is often referred to as a “black box”, and is criticized for being non-transparent and having (for humans) untraceable predictions.

### 5.3 What defines a good binner?

Perhaps the hardest question to answer is how to evaluate a binning tool. There are several questions that come to mind such as, “Are more complete genomes with less precision or less complete genomes with more precision best?” or “When binning multiple samples, if the same genome is present in more than one sample, should it be in a single bin or a bin per sample?”. While the answers to these questions are not completely agreed upon across evaluation tools and benchmarking methods, we will discuss the chosen criteria for this thesis.

For comparing clusters or bins, we chose to use well-developed and throughly-documented clustering performance evaluations. Following prior binning literature [10, 17, 18], clustering metrics such as Rand score, recall, precision, accuracy, and F1 score were chosen.

For evaluating the addition of contig embedding to VAMB, we chose to evaluate the number of near complete (NC) bins, or bins that have at least 0.95 precision and 0.9 recall. Furthermore, we chose to look into high-precision bins of 0.95 precision or higher and minimum recall varying from 0.3 - 0.99.

When it comes to comparing binners, it is important to realize subtle differences in their comparison methods. For example, what does it mean to have 100% recall or precision? For BERT experiments, this means all contigs belonging to the genome must be present in the bin. VAMB on the other hand looks at if there is 100% genome coverage in the bin. By requiring all contigs, the standard for good recall is harder to obtain.

While our evaluation method can be used to compare recall and precision for the methods tested, a direct comparison of these metrics cannot be made when comparing against another binner like VAMB.

### 5.4 Synthetic benchmarking data

We chose to benchmark on synthetic datasets from the Critical Assessment of Metagenome Interpretation (CAMI). CAMI aims to establish standards for benchmarking of metagenomic analysis tools by providing benchmarking datasets [17].

The advantage of using synthetic datasets over real metagenomic datasets is having a “golden standard” or ground truth. In other words, we know which taxonomy group each contig comes from. This allows us to calculate the recall and precision of all the bins against a ground truth rather than a reference database.

The main disadvantage of synthetic data is a lack of realistic sequencing and assembly. Simply put, real metagenomics is “messy” and quality of assemblies are often quite poor. Synthetic data is often assembled too well which means we risk overestimating performance compared to what is feasible with real metagenomic data. Furthermore, a significant difference in our data quality may result in our binner not performing well on a real metagenomic dataset.

The comparison of real metagenomic data against synthetic is somewhat paradoxical as our binners can only be accurately tested on synthetic data, but for the purpose of working well on real data. One way to help fix this is to create more “realistic” datasets. CAMI has recently released new synthetic datasets that are meant to be more realistic. One could hope that training a deep learning model on more realistic datasets will lead to an increased performance for real metagenomic data. Another option is using mock communities, where you have many isolated, cultured bacterial strains with known ratios. This approach provides real sequencing data while also knowing the ground truth, but is limited to cultivable bacteria [20].

### 5.5 Summary of experimental results

The purpose of the masked k-mers task was to create a deeper semantic understanding of a contig as compared to a single extracted contig feature. As the masked k-mers task did not embody the relationship between contigs, the next segment prediction (NSP) was added for the latter part of the BERT model’s training. The NSP task was added in hopes of the model learning the relationship between multiple segments of a contig.

The lack of improvement following 20 epochs of NSP/masked-kmer training, could indicate NSP being an improper task for learning the semantic relationship between segments of a contig. Another possibility could that it was the decreased size of the input rather than training task, causing no significant improvement. As the model takes a maximum input of 512 k-mers, the segments were decreased to a size of 254. The masking task was therefore applied to two 254-segments rather than one 512-segment as previously. This decrease in information, may have hindered the model in learning the required semantic relationships both across segments of contigs, and for each individual segment.

Further experiments are needed in order to conclude if the NSP task or shorter segment length led to a lack of significant improvement in binning performance. This could be done by training a BERT model for 20 epochs on the NSP/masked k-mer task to begin with and compare the results to the BERT model trained only on the masked k-mers task.

As shown in the results, there is a clear trade-off between precision and recall when binning contigs. An increase in precision led to a decrease of recall and vice-versa. Ideally, we want both values to be as high as possible. However, this may not be possible.

Choosing whether you want high precision or recall ultimately comes down to your research question and reason for binning. One may opt for a higher precision clustering algorithm if false negatives are more preferable to false positives. For metagenomic binning, false negatives could be preferred if the goal was to identify if a genome was, with certainty, present in a sample. Alternatively, one may opt for a higher recall clustering algorithm if false positives are more preferable to false negatives. For metagenomic binning, false positives would be preferred if the goal was to discover as many new genomes as possible in a sample.

### 5.6 Is the traditional contig filtering too strict?

The comparison of TNF to DeepBin’s latent space embedding shows that TNF performs better on the longest contigs whereas DeepBin performs equally well across all contig lengths. This is expected as DeepBin is getting a constant 512-segment size regardless of contig length where TNF’s signal is largely dependent on contig length.

As the results provided are only based on a single sample, it is important to consider that every sample and dataset will have a unique distribution of contig lengths. As neither TNF nor DeepBin dominated at all lengths, it was concluded that contig embedding would provide best results as a supplement to TNF rather than replacement.

Typically, prior to binning, short contigs under 2000-2500 bp are filtered out. This strict filtering is needed as the k-mer composition and estimated abundance are highly variable for smaller contigs, meaning the signal from both features gets cancelled out among the noise. Additionally, there is a risk that these small contigs may negatively affect the clustering algorithms performance on longer and “easier” to bin contigs.

The common solution for binners has therefore been strict contig filtering. However, filtering contigs removes useful data that could provide more information about the sample. In fact, for most metagenomic datasets, the majority of contigs will be under 2,000 bp. For example, only 201,253 out of 2,964,257 contigs are kept when filtering the CAMI Oral Dataset for contigs under 2000 so 97% of contigs filtered out!

DeepBin’s ability to form semantic relationships for contigs of all lengths allows for less strict filtering of contigs. The hope is with less strict filtering and more contigs, the reconstruction of higher recall bins is possible.

### 5.7 Further work

Training a BERT model is computationally intensive and costly. This led to a disparity between the needed resources and what were available for this thesis. During training, the model never showed sign of converging and we therefore, hypothesize training the model longer would significantly improve binning results.

Following similar modifications that were taken to improve the original BERT language model [21], we suggest training the model for longer on more datasets, with only the masked k-mers task. Ideally, training the BERT model on more metagenomic datasets could allow the model the ability to better generalize when given new data. Furthermore, these results should be benchmarked with more state-of-the-art binners such as MetaBAT2 [22] and MaxBin2 [23].

As shown in Chapter 2 Methods, for longer contigs, we split them into multiple segments (of length 512) in order to feed them to the BERT model. This means when the contig is embedded it is only based on a random 512-segment, which may or may not have been previously trained on. Future studies should be pursued to provide additional contig information to the model. This could be done by combining every 512-segment embedding for a contig. To do this, the segments should be independently fed to the model and the final representations of each segment should then be averaged together to represent the embedding for the entire contig. An alternative direction would be to increase the BERT model’s maximum sequence length. Recently there’s been several methods that address Transformer input size, notably, Big Bird [24], which shows Transformer-based models working on up to 4096 words. In addition, a recent paper [25] presented a method which allowed training on smaller segments while extending to arbitrary lengths at inference time.

Lastly, future studies could be done to refine the output of the current BERT model. One way would be refining the BERT output with contig abundance as that information is not provided to the model.

## 6 Conclusion

Recent unsupervised deep learning models have demonstrated the ability to capture hidden syntax, grammar, and semantics within DNA sequences [9]. My major contribution is further generalizing these findings to deep bidirectional architectures, allowing the same pre-trained model to successfully tackle the metagenomic binning task.

We introduce contig embedding, a feature which globally captures contextual information for the entire input segment. Results on the CAMI Oral dataset shows incorporating the contig embedding feature is able to improve overall binning results. Specifically, we show that contig embedding is superior to TNF for short contigs in addition to reconstructing more high precision bins for both trained and unseen data. Additionally, DeepBin is the first binner to use contigs as a direct input rather than extracted input features.

We believe that the BERT model has strong potential in assisting biological breakthroughs and state-of-the-art results for genomic tasks through an increased understanding of DNA. We hope that the availability of the code (github.com/ninashenker/deepbin) will inspire the further use of the Transformer architecture for metagenomic binning and for the development of deep learning metagenomic pipelines.

## 7 Supplementary Material

## Footnotes

1 Tetranucleotide frequency is also defined using k-mers of size 4.

2 A single value expressing the significance of the accumulated data

3 Tensor - a mathematical object analogous to but more general than a vector, represented by an array of components that are functions of the coordinates of a space.

4 From a mathematical standpoint, Rand index is related to the accuracy, but is applicable even when unique cluster labels are not used.

5 The F1 score is calculated from the precision and recall of the bins and is the harmonic mean of both measurements.

6 BAM files are generated as output by the short read aligner, Minimap2 [19]

7 768 is the number of features embedded by the BERT model

8 Recalling from methods, near-complete (NC) bins are defined as bins with recall 0.9 or higher and precision 0.95 or higher.

9 With the model input limitation of 512 k-mers, we cannot gain a full global understanding for longer contigs, but comparing only contigs under 512, it would be the case.

